# Selection on sperm size in response to promiscuity and variation in female sperm storage organs

**DOI:** 10.1101/2022.08.05.502910

**Authors:** Emily R. A. Cramer, Zelealem B. Yilma, Jan T. Lifjeld

**Affiliations:** Sex and Evolution Research Group, Natural History Museum, University of Oslo; Carnegie Mellon University in Qatar

**Keywords:** sperm morphology, sperm storage, mate choice, cryptic female choice, sperm length

## Abstract

Sperm cells are exceptionally morphologically diverse across taxa. However, morphology can be quite uniform within species, particularly for species where females copulate with many males per reproductive bout. Strong sexual selection in these promiscuous species is widely hypothesized to reduce intraspecific sperm variation. Conversely, we hypothesize that intraspecific sperm size variation may be maintained by high among-female variation in the size of sperm storage organs, assuming that paternity success improves when sperm are compatible in size with the sperm storage organ. We use individual-based simulations and an analytical model to evaluate how selection on sperm size depends on promiscuity level and variation in sperm storage organ size (hereafter, female preference variation). Simulated species with high promiscuity showed stabilizing selection on sperm when female preference variation was low, and disruptive selection when female preference variation was high, consistent with the analytical model results. With low promiscuity (2-3 mates per female), selection on sperm was stabilizing for all levels of female preference variation in the simulations, contrasting with the analytical model. Promiscuity level, or mate sampling, thus has a strong impact on the selection resulting from female preferences. Further, for species with low promiscuity, disruptive selection on male traits will occur under more limited circumstances than many previous models suggest. Variation in female sperm storage organs likely has strong implications for intraspecific sperm variation in highly promiscuous species, but likely does not explain differences in intraspecific sperm variation for less promiscuous taxa.

## Introduction

Sperm cells have exceptional morphological diversity across species (Pitnick *et al*., 2009). This diversity is partly driven by fertilization environment (internal vs. external; Kahrl *et al*., 2021b), and is also hypothesized to be driven by sexual selection, which can arise when a female copulates with multiple males in a single reproductive bout. With such female promiscuity, sperm from different males may compete to fertilize the egg(s) (Parker, 1970) and/or the female may exert cryptic choice for particular sperm or male characteristics (Eberhard, 1996). How (and whether) such post-copulatory sexual selection processes result in selection on sperm morphology requires more study in most study systems (Lüpold & Pitnick, 2018), but two patterns are quite robust across studies. Specifically, sperm cell morphology co-evolves with the morphology of female sperm storage organs both in comparative studies (Dybas & Dybas 1981; Briskie & Montgomerie 1992; Higginson *et al*. 2012; reviewed in Lüpold & Pitnick 2018) and in experimental evolution studies (e.g., Hosken *et al*. 2001; Miller & Pitnick 2002). These studies suggest that sperm evolve to “fit” sperm storage organs (and/or vice versa) in internally fertilizing species. In addition, among-male variation in sperm length is lower in more promiscuous taxa, suggesting stronger selection for an optimal sperm phenotype (sperm total length: birds, Calhim *et al*. 2007; Lifjeld *et al*. 2010; rodents, Varea-Sánchez *et al*. 2014; and social insects, Fitzpatrick & Baer 2011; flagellum length: sharks, Rowley *et al*. 2019). In this paper, we use simulations and an analytical model to explore how promiscuity level and among-female variability in the sperm storage organs interact in driving selection on sperm.

Female sperm storage organs represent an important selective environment for sperm cells in many species. Correlations between individual males’ proportion of sperm stored and proportion of eggs fertilized can be high, reinforcing the idea that successful interaction with the female is important (Bretman *et al*. 2009; Manier *et al*. 2010; Hemmings & Birkhead 2017; though note that females do not necessarily use stored sperm from all males, e.g. Simmons & Beveridge 2010; Turnell & Shaw 2015). Many factors may impact the successful storage of sperm, including motility as the sperm enter the sperm storage organ (Mendonca *et al*., 2019), mating order (Hellriegel & Bernasconi, 2000; Manier *et al*., 2010; Hemmings & Birkhead, 2017), complex biochemical interactions among ejaculates and with the female (den Boer *et al*., 2010), and genetic compatibility of the male and female (Simmons *et al*. 2006; Gasparini & Pilastro 2011; though genetic compatibility may be assessed in the male rather than directly from the sperm, Løvlie *et al*. 2013). Here we focus on the potential impact of morphological compatibility between the sperm cell and the sperm storage organ, which is suggested by the co-evolution of morphology of sperm and sperm storage organs across taxa (reviewed in Lüpold & Pitnick 2018). There are notable exceptions to the idea of morphological compatibility; for example, García-González and Simmons (2007) find stronger selection for short sperm in females with larger sperm storage organs in the dung beetle *Onthophagus taurus*, so the mechanism we outline here will not be applicable in all systems.

In addition to being important selective environments for sperm, female sperm storage organs likely vary among individuals, following several lines of evidence. First, since genetic variation is a pre-requisite for evolution, the fact that sperm storage organ morphology evolves suggests that it varies (Jennions & Petrie, 1997). Genetic variation in sperm storage organ morphology has also been directly documented (Miller & Pitnick, 2002; Lüpold *et al*., 2013). In addition, environmental and social conditions during development can affect sperm storage organ morphology (Amitin & Pitnick, 2007; Berger *et al*., 2011; Farrow *et al*., 2022). Within-female variation is also known, for example, in birds, where each female has hundreds of sperm storage tubules, whose lengths vary in a gradient across the utero-vaginal junction (where these structures occur), and with stage of the egg-laying cycle (Briskie, 1996).

Thus we hypothesize that females vary in their sperm storage organ morphology, and that the morphological fit between these organs and sperm cells is a mechanism of cryptic female choice, because it biases storage success (and therefore fertilization success) towards well-fitted sperm. We model a scenario where all females have the same preference function, whereby the sperm that best fit their sperm storage organs is more likely to fertilize their eggs. However, females’ preferences (i.e., the sperm trait values that best fit individual females) vary because the preference function is self-referential against a variable morphological trait. This hypothesis is supported by Hemmings et al. (2016), who allowed females to copulate with one male and then compared the morphology of ejaculated cells and of sperm cells that reached the ovum after sperm storage. Re-analysis of their data (see Supplemental file) indicates that the mean sperm length at the egg differed significantly from the mean ejaculated sperm in 10 of 27 females (Table S1). Sperm at the egg was longer than ejaculated sperm for half the females and shorter in the other half, consistent with variable female preferences for sperm size. Furthermore, under this hypothesis, we can expect that males may have different relative fertilization success when they copulate with different females. Several studies do indeed find that the combination of male and female identities (or genetic lines) has a strong impact on fertilization success (Wilson *et al*., 1997; Clark, 2002; Birkhead *et al*., 2004; Bjork *et al*., 2007; Simmons *et al*., 2014; Reinhart *et al*., 2015) (although we note that a combinatorial effect of male and female may also arise due to other processes, for example, variation in copulation duration, Eady and Brown 2017, or sperm swimming speed, Urbach *et al*. 2005; Cramer *et al*. 2014; Cramer *et al*. 2016).

Because we view the fit of sperm and sperm storage organ as a mechanism of cryptic female choice (Lüpold & Pitnick, 2018), we can expect some parallels between this process and mate choice. However, to our knowledge, no theoretical work on mate choice models the conditions most relevant for sperm-female interactions. Specifically, most mate choice models assume that females copulate with a single male in the population, while empirical data show that females often copulate with multiple males, who then share paternity of their offspring (e.g., Gage, 1994; Simmons *et al*., 2007; Simmons & Beveridge, 2010; Turnell & Shaw, 2015; Brouwer & Griffith, 2019; Kahrl *et al*., 2021a). In addition, we assume that females copulate with fewer males than they assess during mate choice, implying that females sample the sperm of relatively few males. The number of sampled partners is known to impact resulting selection strength (Janetos, 1980; Gomulkiewicz, 1991; Muniz & Machado, 2018). Finally, in species where eggs are ovulated in batches, female sperm storage organs have already gathered all the sperm cells that potentially can fertilize the eggs, making cryptic female choice best represented by a simultaneous assessment model. Under a simultaneous assessment strategy, the female evaluates all individuals in a set of potential males before choosing among them. Simultaneous assessment strategies can give different results from other assessment strategies (Janetos, 1980; Jennions & Petrie, 1997; Muniz & Machado, 2018), and to the best of our knowledge, continuous variation in female preferences has not been modeled with simultaneous assessment with a reasonable number of copulation partners (for an internally fertilizing species). See Millan *et al*. (2020) for relevant work with a different assessment model, and Van Doorn *et al*., (2001) and van Doorn & Weissing (2002) for models relevant for broad-cast spawners with high mate sampling. Further work is thus needed to understand how variation in female sperm storage organs impacts selection on sperm.

Here, we use individual-based simulations and an analytical model to investigate how among-female variation in sperm storage organs affects the resulting selective pressure on sperm, and we assess whether this relationship depends on the level of female promiscuity, ie., number of copulation partners. We predict that selection will be stronger with higher promiscuity (Janetos, 1980; Gomulkiewicz, 1991; Muniz & Machado, 2018). We further hypothesize that where female preference is less variable than sperm, there will be stronger stabilizing selection on sperm as female trait variation is further reduced. Conversely, where female preference is more variable than sperm, we predict that there will be stronger disruptive selection on sperm as variation in the female trait increases (Jennions & Petrie, 2000; Van Doorn *et al*., 2001; van Doorn & Weissing, 2002; Weissing *et al*., 2011).

## Methods

### Assumptions

We assume a closed population with an equal sex ratio, where copulations occur randomly with respect to the sperm and preference traits. All eggs are fertilized, so that preferences are selectively neutral for females. This assumption is similar to the “last-chance” option of Janetos (1980), whereby females accept any male rather than not mate.

### Simulation procedure

For each iteration of the simulation, we created a population of 200 individuals of each sex, breeding for one season. Each female produced one set of 25 offspring. This value was chosen to enable us to exploring a relatively high number of copulation partners, while still allowing a substantial probability for each partner to sire offspring. Males are assigned a sperm trait from a normal distribution with mean 0 and SD = 1. Females are assigned a preference (i.e., sperm storage organ size) on the same scale, such that the fit between sperm and preference is best when the trait values are equal. We varied population-level SD in female preference (values of 0.5, 1, 1.5, and 2; comparable to the variation explored by Millan *et al*. 2020), but, for simplicity, the population mean preference was always equal to the sperm preference mean.

All individuals copulated with members of the opposite sex 1, 2, 3, 5, 10, or 25 times. Detailed information on number of copulation partners is poorly known for many species, and is often inferred from genotyping stored sperm in the female or determining paternity of offspring. Empirical data thus provides a minimum estimate of number of individual partners (Cramer *et al*., 2020a). For many species, an average number of copulation partners less than 5 appears realistic (Gage, 1994; Brommer *et al*., 2007, 2010; Simmons *et al*., 2007; Simmons & Beveridge, 2010; Turnell & Shaw, 2015; Cramer *et al*., 2020a; Kahrl *et al*., 2021a), though in eusocial insects the average can be over 50 (Tarpy *et al*., 2004). The values we chose to investigate were also informed by the expectation that selection strength should asymptote with > about 10 copulation partners (Gomulkiewicz, 1991; Muniz & Machado, 2018). We include 1 copulation partner to confirm the expectation of no selection on sperm under this condition. Copulation partners were assigned randomly by shuffling the list of individual identities for each copulation event. This could result in in a pair of individuals copulating with each other more than once. Since that presumably occurs in nature and represents a limited proportion of copulations, we do not control for such repeated copulations in statistical analysis.

Following copulation, the fertilizing sperm for each egg was determined using R’s *sample* function. Weighting the *sample* function requires positive, non-zero values; it then sums all individuals’ weight values, and the probability that an individual is drawn is proportional to its contribution to the sum of the weight values across all individuals. Thus, a male’s success depends on his relative weight (i.e., fit) for the female preference compared to the other copulation partners, not his absolute fit. To calculate absolute fit, we modeled the fit quality as a Gaussian function which attains its maximum when the male’s trait value, *y*, matches the female’s preference, *x*. The parameter *σ*_*U*_, akin to standard deviation, controls the strength of the preference (sensu Millan *et al*., 2020). For the sake of simplicity, we use a value of *σ*_*U*_=1 in all simulations. We therefore calculated the fit score between the sperm size, *y*, and the female preference, *x*, as:

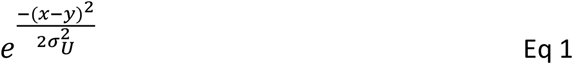

This equation represents the preference function used by all females. After calculating the fit for all copulation partners, we assigned fertilization by drawing male identities from a list of the individual female’s copulation partners, weighted according to the fit scores.

After counting all offspring sired for each male, the selection gradient on the sperm trait was calculated. To do so, reproductive success was standardized by dividing by the population mean reproductive success. Sperm trait values were standardized to have a population mean of 0 and standard deviation of 1 (following Lande & Arnold, 1983). Standardized reproductive success was then regressed on the standardized sperm trait, including both a linear and a quadratic term (Lande & Arnold, 1983). Negative values of the quadratic term indicate stabilizing selection, and positive values indicate disruptive selection. We extracted the quadratic selection gradient parameter from each replicate population.

After performing 1000 replicate populations with the same set of conditions, we compared how the quadratic selection gradient changed with the treatments (variation in female preference and number of copulation partners). To facilitate interpretation, we treat each predictor as categorical rather than continuous. Following the logic outlined in White *et al*. (2014), we rely on effect size estimates rather than p-values in interpreting our results (since simulations can make sample size be arbitrarily high and p-values correspondingly low). Following Richardson (2011), we use η^2^ as the effect size estimate, with values of 0.1, 0.25, and 0.5 considered small, medium, and large, respectively. These were calculated via sjstats (Lüdecke, 2021). We further directly calculated the 95% confidence limits on each simulation condition, as the 2.5% and 97.5% quantiles of the observed values.

All simulations and statistics were performed in R using base functions and the tidyverse package (Wickham *et al*., 2019). We ran a modified set of simulations to assess the impact of sharing paternity (Table S2) and of having larger clutch size relative to copulation partner count in an open population (Table S3). Overall patterns were highly similar.

### Analytical model

Among-female variation in female preference had strong impacts on the shape of selection (see Results), which depended also on the number of copulation partners. To better understand when disruptive or stabilizing selection should be expected when the female could sample all males, we used an analytical model that parallels the simulation. Similar to the simulations, among-female variation in preference is modeled as normally distributed with mean of 0 and standard deviation *σ*_*F*_. Among-male variation in sperm is modeled as normally distributed with mean 0 and standard deviation *σ*_*M*_. The probability density function of the female preference across all females is then

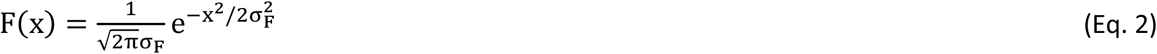

and the probability density function for the sperm trait across all males is

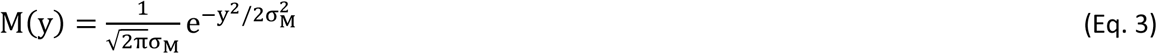

Denoting the preference function as U(x,y), the probability distribution function of fertilization success for all males with trait value *y*, given a female with preference *x*, can be expressed as

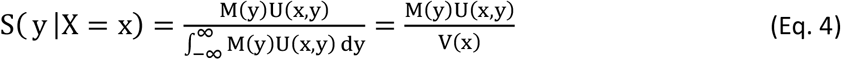

Intuitively, the denominator, V(*x*), can be thought of as the total of the female’s fit scores across all males in the population, and the numerator expresses the contribution of males with trait value y to the total of the female’s fit scores. This is analogous to the *sample* function if all males were sampled.

Fertilization success for all males with trait value *y* can be calculated as the integral of their fertilization success across all females:

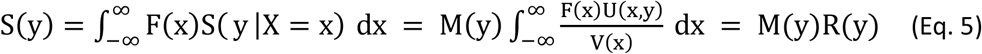

where

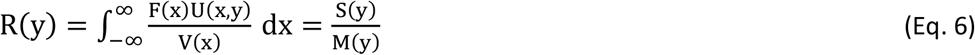

gives the fertilization success of males with value *y*, relative to their representation in the population.

In the simulations, we assumed that U(*x,y*) was given by Eq. 1. Under this condition, we can explicitly calculate the function R(*y*). By substituting Eq 1-3 into the more general form equations 4-6, we have

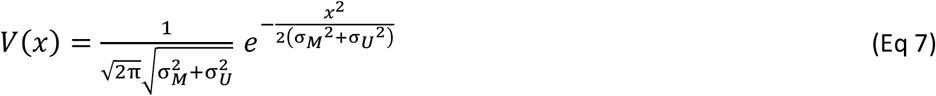

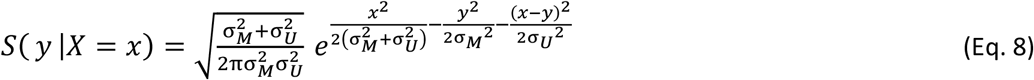

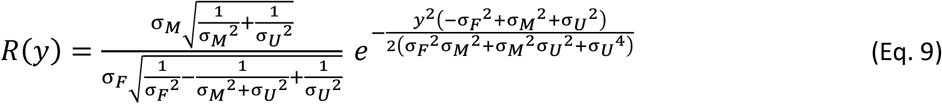

We can re-write Eq. 9 as follows:

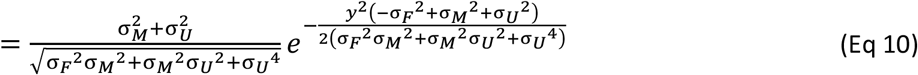

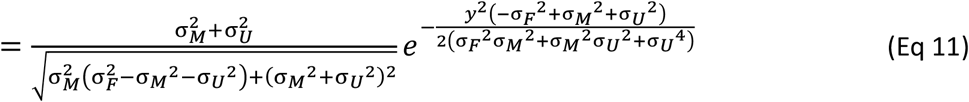

This model is similar to models used by several authors (e.g., Lande, 1981; Dieckmann & Doebeli, 1999), but those authors did not explicitly describe conditions predicting stabilizing and disruptive selection.

## Results

### Simulations

The value of the quadratic selection gradient term depended on among-female variation in preference (F_3,27972_ = 74073, η^2^ = 0.54), number of copulation partners (F_6, 27972_ = 11557, η^2^= 0.17), and the interaction between the two variables (F_18, 27972_ = 5169, η^2^= 0.23; Fig. 1, Fig. 2). Quadratic selection estimates changed dramatically with variation in female preference when the number of copulation partners was high. However, when females copulated only twice, stabilizing selection was only slightly weakened by higher variation in female preference (Table 1). There was no selection on sperm when the female copulated with only one male (Table 1, Fig 2). Patterns were similar in the additional simulation conditions tested (e.g., larger number of offspring per female; see supplementary information).

**Table 1.** Median and 95% quantile limits on the quadratic term from selection gradients

| N copulation partners | Among-female SD in preference |  |  |  |
| --- | --- | --- | --- | --- |
|  | 0.5 | 1 | 1.5 | 2 |
| 1 | 0.00 (-0.00, 0.00) | 0.00 (-0.00, 0.00) | 0.00 (-0.00, 0.00) | 0.00 (-0.00, 0.00) |
| 2 | -0.16 (-0.21, -0.12) | -0.13 (-0.18, -0.09) | -0.11 (-0.16, -0.06) | -0.09 (-0.14, -0.03) |
| 3 | -0.19 (-0.24, -0.14) | -0.15 (-0.20, -0.10) | -0.10 (-0.15, -0.05) | -0.06 (-0.12, 0.00) |
| 5 | -0.21 (-0.26, -0.16) | -0.14 (-0.19, -0.10) | -0.07 (-0.13, -0.02) | -0.01 (-0.07, 0.06) |
| 10 | -0.22 (-0.27, -0.17) | -0.14 (-0.18, -0.09) | -0.03 (-0.09, 0.03) | 0.07 (-0.00, 0.15) |
| 15 | -0.22 (-0.28, -0.17) | -0.13 (-0.18, -0.09) | -0.02 (-0.07, 0.05) | 0.11 (0.03, 0.19) |
| 25 | -0.22 (-0.28, -0.17) | -0.13 (-0.18, -0.09) | -0.00 (-0.05, 0.06) | 0.14 (0.07, 0.23) |

**Figure 1.**
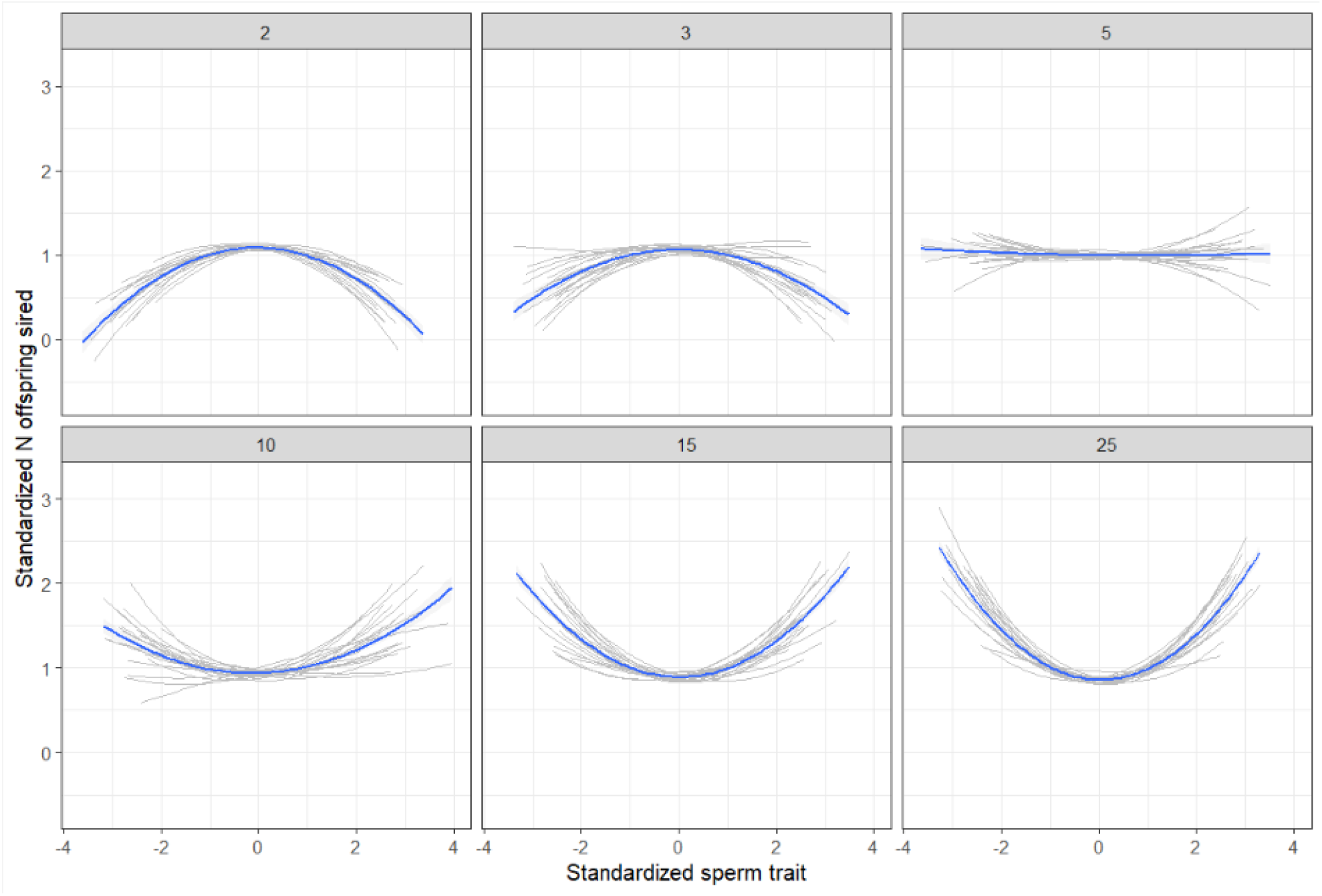
Examples of selection gradients on the sperm trait for 20 randomly selected populations with the SD for among-female variation in preference = 2. Each panel shows a different level of number of copulation partners. The male trait had a standard deviation of 1 in all treatments. Grey lines show 20 randomly selected individual populations and the blue line shows the overall pattern.

**Figure 2.**
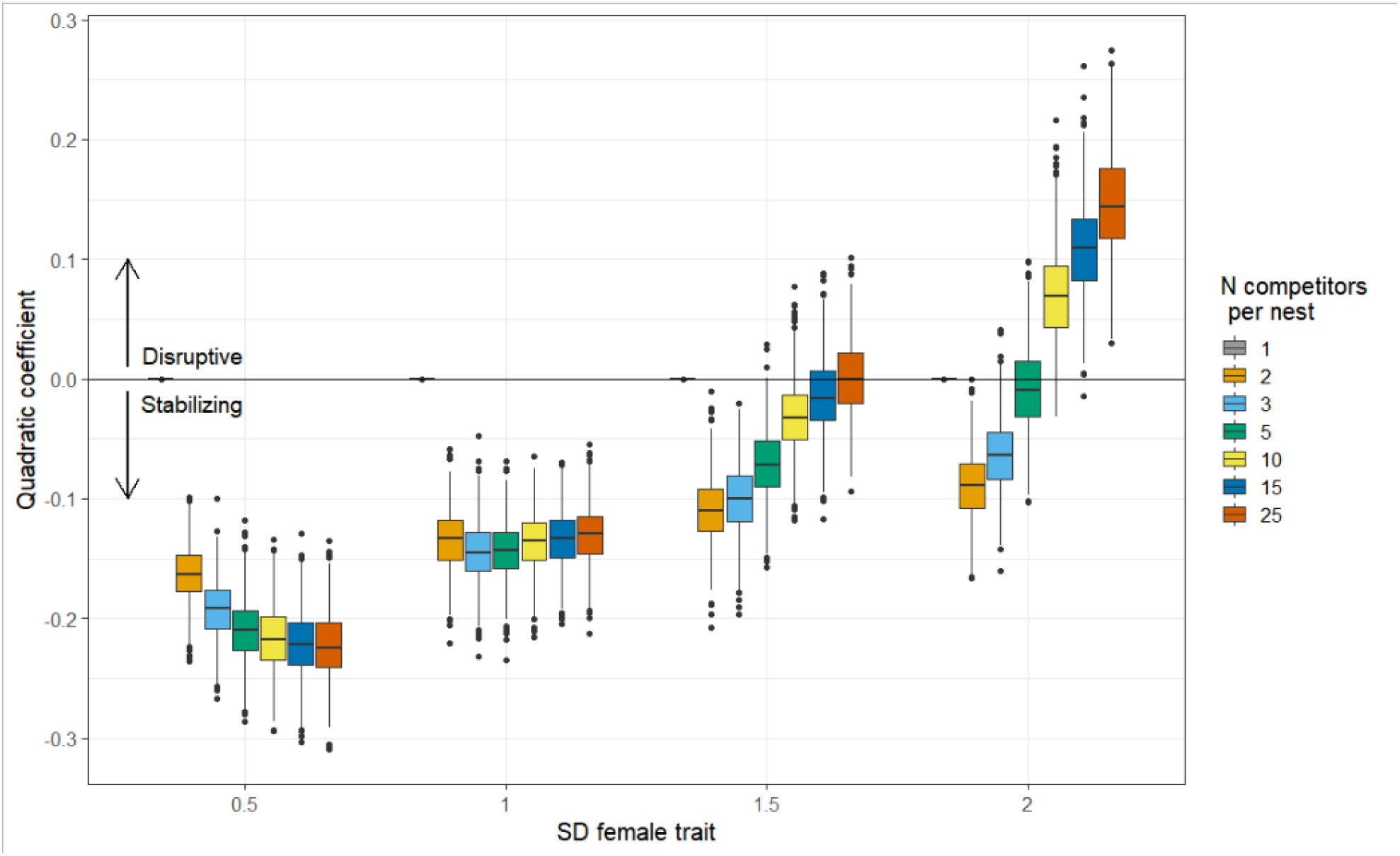
Values of the quadratic selection gradient for all simulation conditions. Colors indicate the number of copulation partners. A value of 0 (heavy horizontal line) indicates no quadratic selection; negative values indicate stabilizing selection; and positive values indicate disruptive selection. Results include 1000 replicates per simulation condition.

With female preference SD 0.5, there was stabilizing selection on sperm (negative quadratic selection coefficients, with 95% quantiles not overlapping 0), and the median strength of stabilizing selection increased with the number of copulation partners between 2 and 5, but did not increase between 5 and 25 (Table 1, Figure 2). With female preference SD 1, i.e., equal to male trait SD, selection on sperm was again stabilizing, but here there was little variation depending on number of copulation partners. When the SD of the female preference was 1.5, selection was stabilizing with 2, 3, or 5 copulation partners, and there was no selection with 10, 15 or 25 copulation partners (95% quantiles include 0). When the SD of the female preference was 2, selection was stabilizing with 2 copulation partners; there was no selection with 3, 5, or 10 partners, and there was disruptive selection with 15 or 25 partners (95% quantiles not overlapping 0, positive quadratic selection coefficients). The impact of increasing the number of copulation partners appeared to asymptote here, as the difference in median disruptive selection between 5 and 15 partners was 0.1, while the difference in median disruptive selection between 15 and 25 partners was only 0.03 (Table 1).

### Analytical model

From Eq 11, we see that the shape and intercept of R(y) is determined by the expression −σ_*F*_ ^2^ + σ_*M*_ ^2^ + σ_*U*_ ^2^. In particular, if σ_*F*_ ^2^ < σ_*M*_ ^2^ + σ_*U*_ ^2^, R(y) is bell-shaped and has R(0) > 1 as its maximum value, indicating that males with average trait values gain greater fertilization success than would be expected given their frequency in the population. This implies stabilizing selection. If σ_*F*_ ^2^ > σ_*M*_ ^2^ + σ_*U*_ ^2^, R(y) is U-shaped and R(0) < 1, indicating that males with average trait values gain less fertlization success than expected and implying disruptive selection. No selection is expected where σ_*F*_ ^2^ = σ_*M*_ ^2^ + σ_*U*_ ^2^ as this results in the constant function R(y) = 1. We evaluated whether this result agreed with model results by arbitrarily choosing several sets of values for the three variances that should give no quadratic selection (see supplementary materials, Table S4).

## Discussion

Stabilizing selection on the sperm trait when there is less variation in female preference than in sperm is intuitive: all females share a preference for sperm with a phenotype close to the male population mean. Disruptive selection when female preference is more variable than sperm is similarly intuitive: many females have preferences outside the sperm trait distribution, thus the most extreme males in the population obtain high fertilization success after copulating with a matching female (Van Doorn *et al*., 2001; van Doorn & Weissing, 2002; Millan *et al*., 2020). Our analytical model indicates that the change from stabilizing to disruptive selection should occur when the among-female preference variance is greater than the sum of the among-male variance in sperm traits and the variance parameter in the female preference function (Eq 1, which we do not vary in this simulation). Our simulation results, however, show stabilizing selection under more conditions than expected, also compared to previous models where females sampled a large subset of males (Millan *et al*., 2020). Specifically, when the number of copulation partners is low, stabilizing selection can occur even with high among-female variation in preference. We suggest that this stabilizing selection occurs because males with relatively extreme sperm values are relatively unlikely to copulate with females with a matching preference, and their fertilization advantage when they do achieve these matching copulations is insufficient to offset the rarity of the copulations. The importance of sampling number is also evident in the empirical literature, where mating preferences are expressed more strongly in studies where individuals can choose among two mating options, compared to studies where individuals have a single option and can mate or not (Dougherty & Shuker, 2015).

### Implications for sperm evolution

This study shows that variation in cryptic female preferences and variation in number of copulation partners each can have a strong impact on the strength, or even shape, of selection on sperm morphology. Perhaps surprisingly, there are conditions where number of copulation partners does not impact the strength of selection (i.e., where sperm and preference variation are equal), and there are conditions where no selection on sperm is expected even when there are a large number of copulations. It is difficult to know which combination of variables is likely to be most biologically relevant, since variation in female genital morphology is under-studied relative to male genital traits (Ah-King *et al*., 2014), and copulation behavior is difficult to observe in the wild. However, we can draw some generalizations. With low levels of promiscuity (2-3 copulations), selection is expected to be stabilizing, and it is similar across levels of variation in female preference. In contrast, with high numbers of copulations (10-25), selection on sperm is stabilizing, null, or disruptive, depending on the level of variation in the female preference.

#### Low to moderate promiscuity systems

For many species, we suspect that the number of copulation partners is low enough that stabilizing selection is broadly expected. Inferences of copulation rate from paternity patterns suggest in socially monogamous passerine birds that females on average copulate with fewer than 3 males (Brommer *et al*., 2007, 2010; Cramer *et al*., 2020a). Genotyping remnants of stored sperm in the female reproductive tract indicates that mean number of mates is between 2 and 6 for several invertebrates (including butterflies, crickets, and beetles; Gage, 1994; Simmons *et al*., 2007; Simmons & Beveridge, 2010; Turnell & Shaw, 2015). In such species, assuming heritability of sperm morphology (reviewed by Edme *et al*., 2019), the sperm-female fit function modeled here would then often be expected to erode variation in sperm morphology over time. Why, then, are sperm cells still variable, and why does the level of variability correlate with promiscuity rates?

Stabilizing selection imposed by the need to fit the female’s sperm storage organs may be countered by diverse other selective pressures. For example, different sperm morphology may confer a fertilization advantage depending on whether the sperm are the first-inseminated (ie., in a defensive position relative to competitors) or are later-inseminated (in an offensive role) (Clark *et al*., 1995; Calhim *et al*., 2011). The most advantageous sperm morphology may also depend on the phenotype of the male himself (Ålund *et al*., 2018). Sperm morphology may correlate with other ejaculate traits that are also under selection, such as sperm number and sperm swimming speed, resulting in complex multivariate selection patterns (Snook, 2005; Fitzpatrick *et al*., 2012; Lüpold *et al*., 2012b). Sperm morphology may correlate with pre-copulatory traits under selection (e.g. Simmons *et al*., 2017), creating indirect selection on sperm morphology (Cramer, 2021). Finally, selection for genetically compatible sperm (Simmons *et al*., 2006; Fossøy *et al*., 2008; Bretman *et al*., 2009; Gasparini & Pilastro, 2011; Rekdal *et al*., 2019) is expected to be independent of sperm morphology, since it depends on the genotypes of the male and female. As these examples show, it is most appropriate to consider the sperm-sperm storage organ fit as one component of a complex selective landscape.

At an ontogenetic level, variation in sperm morphology may arise due to various environmental factors, including but not limited to age (e.g., Cramer *et al*., 2020b), seasonal changes in sperm morphology (Lüpold *et al*., 2012a; Cramer *et al*., 2013; Edme *et al*., 2019), larval rearing conditions and timing (Vermeulen *et al*., 2009), differences in the social environment as an adult (Immler *et al*., 2010; Rojas Mora *et al*., 2018), and condition-dependence of sperm phenotypes (which has been documented in some studies but is not generally expected; Macartney *et al*., 2019).

Persistence of variation in sperm morphology may also depend on the genetic and genomic underpinnings of the trait. In zebra finches, for example, a genomic inversion on the sex chromosome allows many loci to act as a super gene influencing sperm morphology (Kim *et al*., 2017), and this supergene shows heterozygote advantage that could sustain genetic variation over time (Knief *et al*., 2017). Maternal genetic effects on sperm traits have been found in several studies (e.g., Ward, 2000; Morrow & Gage, 2001; Froman *et al*., 2002). If the genes causing these maternal effects are X-linked or on the mitochondria, they may be protected to some extent from selection acting on the sperm phenotype (Gemmell *et al*., 2004). Genetic underpinnings of sperm morphology are poorly known for most species, although substantial heritability of sperm morphology indicates strong genetic effects (reviewed in Edme *et al*., 2019). However, heritability is less directly relevant to how quickly a trait is expected to evolve in response to selection than is evolvability (Hansen *et al*., 2011). Evolvability for sperm morphological traits is comparable to values for other linear trait measurements (median 0.1% for linear traits in Hansen *et al*., 2011; range for total sperm length 0.02% - 0.26% in Edme *et al*., 2019, recalculated from CV_A_ to I_A_ for comparability to Hansen *et al*., 2011).

The above examples may help to explain why sperm remain variable despite stabilizing selection, but they do not immediately explain the among-species correlation between promiscuity level and intraspecific sperm morphological variation. Here, at least for social monogamy with extra-pair paternity, we argue that the among-species pattern is likely driven by the proportion of non-promiscuous females in the population. Most socially monogamous species with extra-pair paternity probably include some females that copulate only with their social mate, for example due to successful mate-guarding by that male (e.g., Chuang-Dobbs *et al*., 2001; Brylawski & Whittingham, 2004; Johnsen *et al*., 2008). When a female copulates with only one male and his sperm fertilize all her eggs, she exerts no selection on his sperm morphology. If social mates also copulate much more frequently than extra-pair mates, such that the social male has many more sperm in competition with extra-pair males, selection in the morphology of the social male’s sperm may also be weakened. If we assume that monogamous females exert no selection on sperm and females copulating with 2-3 males exert stabilizing selection (as indicated in the model), then the total strength of stabilizing selection should depend on the proportion of monogamous versus promiscuous females. Assuming that the proportion of monogamous females is lower, or that social males contribute a smaller proportion of sperm, in species with higher extra-pair paternity rates, we then can expect stronger overall stabilizing selection on sperm in those species. Strong stabilizing selection due to high proportions of females obtaining 2-3 copulation partners may result in faster evolution of sperm in these lineages (as seen in Rowe *et al*., 2015), if mean preferences become different from mean sperm traits, for example due to genetic drift.

#### High promiscuity systems

For some groups, for example some eusocial insects, the number of copulation partners can be quite high (Tarpy *et al*., 2004). Here, we expect the shape of selection to depend strongly on the degree of variation in female preference, ranging from stabilizing to disruptive selection. Assuming similar levels of variation in female preference within these taxa, increases in the number of mating partners from an already-high baseline may not result in strong increases in selection strength, due to asymptotic effects. Perhaps in contrast to these expectations, Fitzpatrick & Baer (2011) found a between-species correlation between intraspecific variation in sperm morphology and promiscuity level in eusocial insects, even when excluding monogamous species. However, only five species in one genus (*Apis*) had more than 10 copulation partners in their dataset, limiting the power to test whether promiscuity level correlates with sperm morphological variation only within highly promiscuous species. Our simulation results suggest that selection on sperm will be highly dependent on the degree of variation in the female sperm storage organs in such taxa, although the myriad other factors influencing sperm variation discussed above may also be at play in high-promiscuity systems. The combination of high promiscuity and high variation in female sperm storage organs creates an expectation of disruptive sexual selection, which in turn can play a role in the splitting of lineages to form separate species (Lande, 1981; van Doorn & Weissing, 2002; Weissing *et al*., 2011; see also Van Doorn *et al*., 2001; Howard *et al*., 2009).

### Implications for previous work on mate choice

Our observation that limited mate sampling causes stabilizing selection even with substantial among-female preference variation has important implications for interpreting previous models of sympatric speciation. Previous models have highlighted a broad female preference distribution as a key element in generating disruptive selection on male traits as one step that can lead to sympatric speciation (Higashi *et al*., 1999; Van Doorn *et al*., 2001; van Doorn & Weissing, 2002; Weissing *et al*., 2011). Our results suggest that disruptive selection will occur under more limited circumstances than was previously appreciated, as females generally are expected to be somewhat limited in the number of males they can sample (Jennions & Petrie, 1997). We thus support Servedio & Boughman (2017)’s assertion that novel insights may be obtained in the sympatric speciation literature by further exploring closed-ended preference functions and limited female searches, similar to what we have simulated here.

As expected from previous models (Janetos, 1980; Gomulkiewicz, 1991; Muniz & Machado, 2018), increasing the number of partners generally increased the strength of selection, and the relationships was asymptotic, with the asymptote reached at a lower value in the conditions with stabilizing selection than the conditions with disruptive selection. We further find that selection is generally weaker when paternity is shared within each batch of offspring, compared to when the best-fit male sires all offspring (Table S2). Models of mate choice should therefore use realistic values for number of males sampled and number of males succeeding (in copulating or fertilizing) to obtain the most biologically relevant results.

#### Conclusions

Despite broad interest in sperm morphology, relatively few studies have evaluated selection on sperm morphology in the wild (Lüpold & Pitnick, 2018), and even fewer have evaluated the presence and effect of variation in female sperm storage organs. Under our model where the sperm storage organs bias paternity success towards sperm of a similar size, the level of variation in the female sperm storage organs determines whether selection on sperm is stabilizing or disruptive for highly promiscuous species, whereas selection is stabilizing for species with only 2-3 copulations per female.

## Acknowledgements

Emma Whittington, Thore Koppetsch, Arild Johnsen, and others in SERG provided useful discussion. This study was funded by the Norwegian Research Council, project number 301592.

