## Supplementary Text and tables for "Selection on sperm size in response to promiscuity and variation in female sperm storage organs"

Supplementary information for "Selection on sperm size in response to promiscuity and variation in female sperm storage organs"

*Reanalysis of the data of Hemmings et al. (2016)*

For each male separately, we performed a t-test assuming unequal variances to compare sperm measured from faecal samples (representing ejaculated sperm) and from the perivitelline layer (PVL) of his mate's eggs. Data were downloaded from Dryad on 15 March 2022. Results are provided in Table S1.

Table S1. Summary of t-test results comparing faecal and PVL sperm. Bird identity is the same as listed in Hemmings et al.'s data file. The table is sorted in order of the difference between faecal and PVL sperm (negative scores indicate sperm were shorter in the faecal sample). Significant tests are in bold (no correction for multiple testing).

| Bird Identity | Difference (µm) in sperm total length between faecal and PVL sperm | t-test results |
|---|---|---|
| **1** | **-4.36** | **$t_{10.7}$ = -3.90, p = 0.00** |
| **8** | **-3.70** | **$t_{9.8}$ = -2.61, p = 0.03** |
| **11** | **-2.32** | **$t_{11.2}$ = -3.21, p = 0.01** |
| **3** | **-2.02** | **$t_{11.7}$ = -2.95, p = 0.01** |
| 18 | -1.84 | $t_{10.3}$ = -2.04, p = 0.07 |
| 19 | -1.76 | $t_{10.3}$ = -1.62, p = 0.13 |
| **4** | **-1.67** | **$t_{14.5}$ = -2.42, p = 0.03** |
| 10 | -0.98 | $t_{10.0}$ = -2.09, p = 0.06 |
| 25 | -0.89 | $t_{12.8}$ = -1.35, p = 0.20 |
| 16 | -0.80 | $t_{17.3}$ = -2.00, p = 0.06 |
| 24 | -0.70 | $t_{17.1}$ = -1.45, p = 0.16 |
| 5 | -0.52 | $t_{17.3}$ = -0.77, p = 0.45 |
| 22 | -0.39 | $t_{15.0}$ = -1.33, p = 0.20 |
| 20 | 0.12 | $t_{16.4}$ = 0.47, p = 0.64 |
| 23 | 0.33 | $t_{19.6}$ = 0.32, p = 0.75 |
| 14 | 0.33 | $t_{18.5}$ = 0.59, p = 0.56 |
| 26 | 0.59 | $t_{36.6}$ = 1.61, p = 0.11 |
| 12 | 0.60 | $t_{15.4}$ = 0.70, p = 0.49 |
| **9** | **0.76** | **$t_{14.6}$ = 2.42, p = 0.03** |
| 6 | 0.85 | $t_{19.8}$ = 1.31, p = 0.20 |
| 2 | 1.31 | $t_{11.1}$ = 1.29, p = 0.22 |
| **21** | **1.34** | **$t_{10.8}$ = 3.08, p = 0.01** |
| 15 | 1.39 | $t_{15.5}$ = 1.66, p = 0.12 |
| 13 | 1.42 | $t_{9.6}$ = 1.36, p = 0.20 |
| **7** | **1.51** | **$t_{12.6}$ = 3.66, p = 0.00** |

| | | |
|---|---|---|
| **27** | **1.84** | **$t_{10.7}$ = 3.12, p = 0.01** |
| **17** | **2.52** | **$t_{10.8}$ = 2.81, p = 0.02** |

*Additional simulation conditions*

We ran additional simulation conditions to verify some of our results.

First, in order to evaluate whether shared paternity impacted results compared to most mate choice models where males do not share paternity because females copulate only once, we altered the main model simulation so that the best-fit copulation partner sired all offspring. Here we ran 1000 iterations, with 2, 10, and 25 copulation partners, with female preference SD of 0.5, 1.5, and 2.

We found that sharing paternity weakened selection strength relative to iterations where the best-fit sperm fertilized all eggs (Table S2, Figure S1). However, the main patterns are robust, with selection being either stabilizing or disruptive with high promiscuity, depending on the level of variation in the female preference. When females copulated with only two males, selection was always stabilizing.

Table S2. 95% quantiles for simulations on closed populations where paternity was either shared among males depending on the sperm-sperm storage organ fit ("shared"; these are the main results, re-printed here for ease) or where all paternity was assigned to the single copulation partner with the best-fitting sperm ("unshared"). 1000 iterations were run with 25 offspring per breeding event for the latter.

| N males competing | Is paternity shared? | Female preference SD 0.5 | 1.5 | 2 |
|---|---|---|---|---|
| | shared | -0.16 (-0.21, -0.12) | -0.11 (-0.16, -0.06) | -0.09 (-0.14, -0.03) |
| 2 | unshared | -0.27 (-0.35, -0.20) | -0.14 (-0.21, -0.07) | -0.11 (-0.17, -0.04) |
| | shared | -0.22 (-0.27, -0.17) | -0.03 (-0.09, 0.03) | 0.07 (-0.00, 0.15) |
| 10 | unshared | -0.39 (-0.52, -0.26) | 0.09 (-0.01, 0.22) | 0.25 (0.13, 0.41) |
| | shared | -0.22 (-0.28, -0.17) | -0.00 (-0.05, 0.06) | 0.14 (0.07, 0.23) |
| 25 | unshared | -0.39 (-0.52, -0.28) | 0.27 (0.12, 0.42) | 0.52 (0.34, 0.70) |

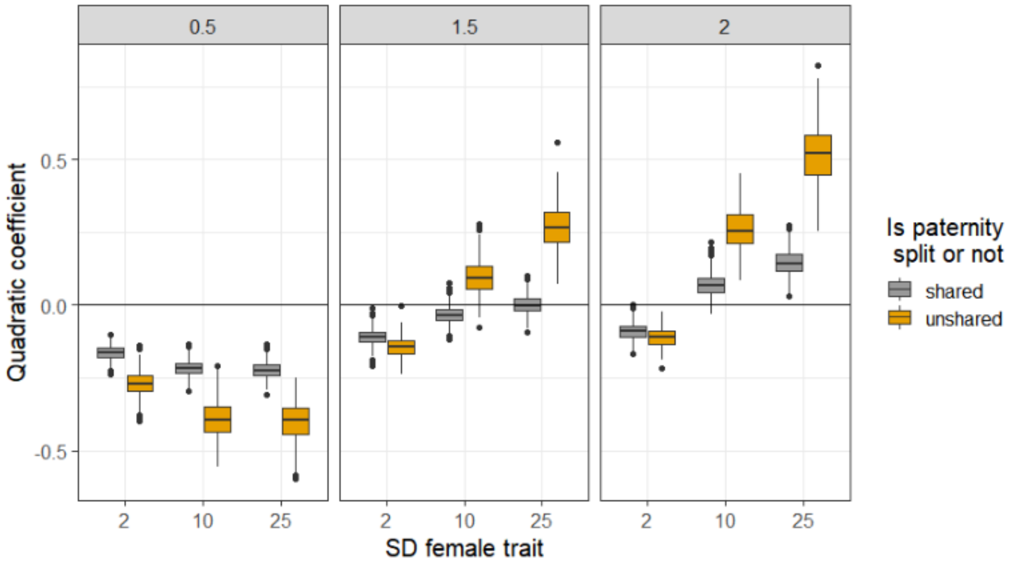

Figure S1. Boxplots showing interquartile range for simulations where paternity was either shared among males depending on the sperm-sperm storage organ fit (“shared”; these are the main results, re-printed here for ease) or where all paternity was assigned to the single copulation partner with the best-fitting sperm (“unshared”).

In addition, we wanted to evaluate the impact of male and female promiscuity separately. In the main simulations, for efficiency and due to the population being a closed population, we coded the simulations such that all individuals of both sexes had the same number of copulation partners. Here, we instead generated populations of 1000 males and 1000 females as described in the main text. We *a priori* chose 200 males to be focal subjects. Each focal male then drew a number of female copulation partners at random without replacement. Each of these females in turn drew a number of copulation partners from the remaining male pool (without replacement). Thus there were no repeated copulations between pairs of individuals within the dataset, and the number of male and female copulation partners could be altered independently. The same females may have been drawn by different focal males, but this was treated as an independent breeding attempt for that female (i.e., she drew another set of copulation partners for that breeding attempt). If a focal male acted as a competitor against other focal males, his paternity in those nests was not included in the analyses. In this simulation, we increased the number of offspring per set to 100, to evaluate whether the asymptoting of selection with increasing number of copulation partners could perhaps have been due to the limited number of offspring relative to copulation partners.

Because this simulation approach was more computationally demanding, we used only a subset of conditions evaluated in the main paper, and we ran only 50 iterations of the simulation. Specifically, we assessed 2, 10, and 25 partners each for males and for females, with female preference SD of 0.5, 1.5, and 2.

Variation in the number of females a focal male copulated with did not impact parameter estimates (Table S3). Values from this modified simulation were highly similar to values from the main simulation with the same number of competitors within a nest (Table S3). We therefore see no evidence that the simpler coding used in the main simulations caused bias.

Table S3. 95% quantiles for modified simulation where number of male and female copulation partners could vary independently and there were 100 offspring per batch. Results from the main simulation for the same conditions are re-printed here (“main”) for easier comparison.

| N males competing within a breeding attempt | N females each male copulates with | Female preference SD | | |
|---|---|---|---|---|
| | | 0.5 | 1.5 | 2 |
| 2 | main | -0.16 (-0.21, -0.12) | -0.11 (-0.16, -0.06) | -0.09 (-0.14, -0.03) |
| | 2 | -0.19 (-0.24, -0.13) | -0.10 (-0.14, -0.02) | -0.05 (-0.14, 0.08) |
| | 10 | -0.19 (-0.23, -0.15) | -0.10 (-0.17, -0.05) | -0.08 (-0.14, 0.03) |
| | 25 | -0.19 (-0.24, -0.15) | -0.09 (-0.14, -0.03) | -0.07 (-0.13, -0.00) |
| 10 | main | -0.22 (-0.27, -0.17) | -0.03 (-0.09, 0.03) | 0.07 (-0.00, 0.15) |
| | 2 | -0.21 (-0.26, -0.17) | -0.05 (-0.15, 0.12) | 0.05 (-0.11, 0.31) |
| | 10 | -0.21 (-0.25, -0.17) | -0.04 (-0.11, 0.06) | 0.07 (-0.05, 0.19) |
| | 25 | -0.21 (-0.26, -0.15) | -0.02 (-0.10, 0.04) | 0.08 (-0.02, 0.18) |

| | | | | |
|---|---|---|---|---|
| 25 | main | -0.22 (-0.28, -0.17) | -0.00 (-0.05, 0.06) | 0.14 (0.07, 0.23) |
| | 2 | -0.22 (-0.26, -0.15) | -0.02 (-0.11, 0.18) | 0.15 (-0.05, 0.38) |
| | 10 | -0.21 (-0.25, -0.18) | -0.00 (-0.11, 0.10) | 0.14 (0.01, 0.34) |
| | 25 | -0.21 (-0.25, -0.18) | -0.00 (-0.10, 0.10) | 0.14 (-0.01, 0.27) |

*Verifying analytical model predictions*

The analytical model predicts no quadratic selection when where $\sigma_F{}^2 = \sigma_M{}^2 + \sigma_U{}^2$ (where $\sigma_F{}^2$is the variance for the whole population in female preference; $\sigma_M{}^2$ is the variance in sperm length; and $\sigma_U{}^2$is the variance in the preference function for each female, which we did not vary in the main simulations). We arbitrarily chose three sets of values for these three terms that satisfy the equation given above, and ran 100 iterations of the simulation for each set of conditions. Here we allowed each individual to copulate 200 times. (Note that this likely resulted in repeated copulations between individuals rather than each individual copulating with all other individuals). Each female produced 25 offspring. As predicted by the analytical model, quadratic selection was 0 for these conditions (Table S4).

Table S4. Median and 95% quantiles for simulations where the expected value of the quadratic selection coefficient is 0, based on the analytical model (based on 100 iterations).

| $\sigma_F{}^2$ | $\sigma_M{}^2$ | $\sigma_U{}^2$ | Quadatic selection coefficient |
|---|---|---|---|
| $\sqrt{2}$ | 1 | 1 | -0.00 (-0.06, 0.04) |
| 4 | 2 | 2 | -0.01 (-0.06, 0.07) |
| 7 | 4 | 3 | -0.01 (-0.06, 0.06) |
